# High resolution AKT signaling in individual cells

**DOI:** 10.1101/373993

**Authors:** Sean M. Gross, Mark A. Dane, Elmar Bucher, Laura M. Heiser

## Abstract

Cells sense and respond to their environment by activating distinct intracellular signaling pathways, however an individual cell’s ability to faithfully transmit and discriminate environmental signals is thought to be limited. To assess the fidelity of signal transmission in the PI3K-AKT signaling pathway, we first developed an optimized genetically encoded sensor that had an increased dynamic range and reduced variation under basal conditions. We then used this reporter to track responses to varying doses of IGF-I in live cells and found that signaling responses from individual cells overlapped across a wide range of IGF-I doses, suggesting limited transmission accuracy. However, further analysis of individual cell traces revealed that responses were constant over time without stochastic fluctuations. We devised a new information theoretic approach to calculate the channel capacity using variance of the single cell time course data‐‐rather than population-level variance as has been previously used—and predicted that cells were capable of discriminating multiple growth factor doses. We validated these predictions by tracking individual cell responses to multiple IGF-I doses and found that cells can accurately distinguish at least four different IGF-I concentrations, as demonstrated by their distinct responses. Furthermore, we found a similar discriminatory ability to pathway inhibition, as assessed by responses to the PI3K inhibitor alpelisib. Our studies indicate that cells can faithfully transmit an IGF-I input into a down-stream signaling response and that heterogeneous responses result from variation in the input-output relation across the population. These observations reveal the importance of viewing each cell as having its own communication channel and underscore the importance of understanding responses at the single cell level.

## Introduction

Cells sense and respond to their local environment by activating distinct intracellular signaling pathways. These pathways are composed of multi-step relays that use second messengers, protein translocations, and kinase/phosphatase cycles to encode, relay, and decode the extracellular signal into an intracellular response^1^. Given the multi-step nature of cell signaling in both space and time, much remains unknown about exactly how individual cells process extracellular signals into accurate intracellular responses^2-5^. Previous studies have observed that cell signaling is inherently noisy and that individual cells are limited in their detection of extracellular stimuli^2-4^, which suggests that ensemble averaging is required to decode extracellular signals into accurate signaling responses. However, these prior studies assessed signaling pathways with predominantly pulsatile‐‐rather than sustained‐‐responses, which hinders the ability to quantify and compare individual cell responses across time and may also obscure alternate explanations for heterogeneity.

The IGF-I ‐ PI3K ‐ AKT signaling pathway provides an excellent model to study cell signaling: it uses multiple modalities to encode, relay, and decode extracellular signals and also produces sustained responses. In brief, IGF-I binds to the receptor tyrosine kinase IGF1R to stimulate intracellular kinase activity^6^. This leads to recruitment of protein binding partners and translation of the protein signal into a lipid second messenger by PI3-Kinase and the subsequent phosphorylation and activation of AKT. The protein kinase AKT can then phosphorylate a range of proteins including the transcription factor FoxO1^6-9^. Previously, we developed a fluorescent fusion protein using FoxO1, which can be used as a readout for IGF-I signaling activity^10^, and found that responses to IGF-I were sustained across time but heterogeneous across the population. We surmised that this heterogeneity could arise from various sources, including: noise in the signaling pathway, variability in the reporter and its quantification, non-specificity in the reporter, or cell-to-cell variation in the encoding of the signal.

Transitions from healthy to diseased states frequently involve changes to cell signaling, therefore a deep understanding of the fidelity of signaling in individual cells can provide insights into disease progression and treatment. For example, cancers arise from genetic alterations that drive activation of oncogenic pathways such as the PI3K-AKT pathway^11-13^. Moreover, cancers are frequently marked by substantial intratumoral heterogeneity^14^, indicating the need to quantify signaling at the single cell level. Several small molecule inhibitors have been developed to inhibit PI3K-AKT pathway activity and block cancer growth^15^. While promising, responses to these compounds are variable and resistance inevitably develops^16,17^. One possible explanation for incomplete therapeutic responses is variation in the signaling capability of individual cells^18-20^. Thus, a more complete understanding of responses to pathway-targeted activators and inhibitors at the single cell level will influence how clinical responses are interpreted and could inform mechanisms of drug resistance.

Here we report the development of an improved FoxO1 reporter to monitor AKT activity in individual cells and use it to assess growth factor signaling responses. In genetically transformed HeLa cells, we observed heterogeneous responses to IGF-I that were consistent with individual cells having different input-output relationships. To account for differences in the input-output relation across the population, we calculated the channel capacity—which provides an assessment of signaling fidelity—by measuring variance in individual cells over a time window when signaling responses had stabilized. This new approach nearly doubled the predicted number of IGF-I doses a cell could accurately discriminate. We then experimentally validated these predictions by treating cells with multiple IGF-I doses and showed that individual cells could accurately discriminate at least four IGF-I concentrations. Furthermore, we found that signaling fidelity extends to pathway inhibition such that the level of activation is largely correlated with the level of inhibition. We conclude that single cell responses to submaximal doses of IGF-I vary across a population because individual cells have different input-output relationships, and further that accounting for these differences reveals high signaling fidelity in individual cells.

## RESULTS

### Construction of a novel FoxO1 reporter with increased specificity

Signaling responses in the AKT, Erk, NF-kB, TRAIL and Ca^2+^ pathways have been shown to be heterogeneous across the population^2-5, 21^. This heterogeneity is frequently interpreted as arising from noise in signal transmission^2-4^. In previous work, we developed a live-cell reporter to study IGF-I responses at the single cell level: a fluorescently tagged FoxO1 protein^10^, which contains three canonical AKT phosphorylation sites that regulate protein localization to the nucleus or cytoplasm depending on their phosphorylation status^9, 22^ (Fig. 1A). We found that the relative fraction of the reporter in the nucleus was variable across a population of HeLa cells cultured in serum free conditions^21^ and speculated that this variation resulted from phosphorylation on other non-AKT phosphorylation sites. In order to assess signaling noise in the IGF-PI3K-AKT pathway, we first sought to reduce basal variation in our readout for pathway activity.

**Fig. 1.**
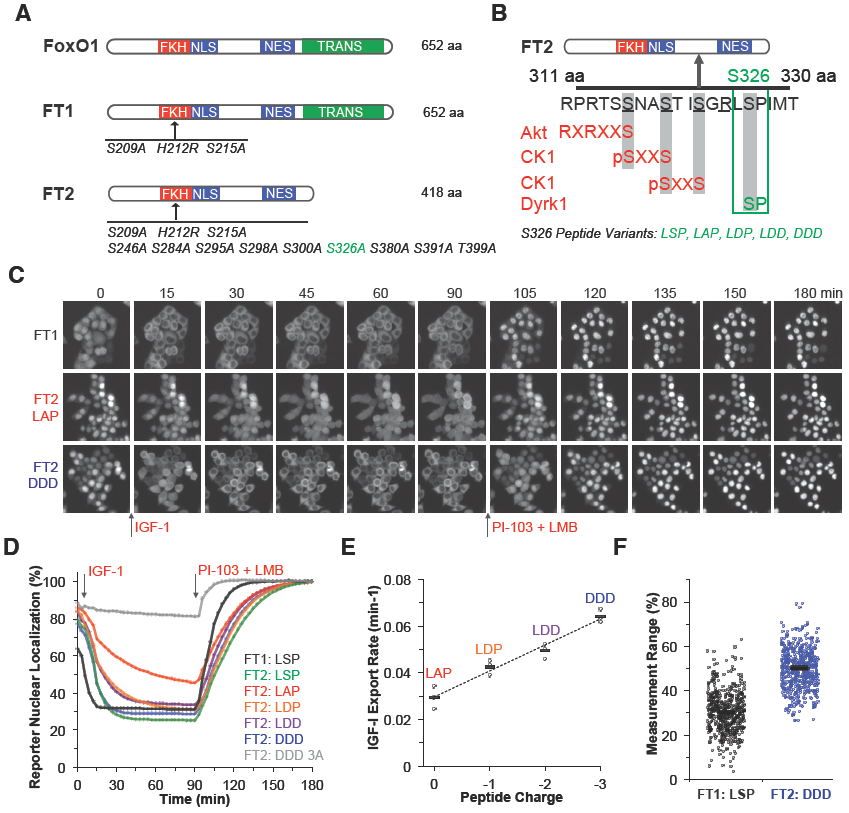
Characterization of an improved FoxO1 reporter. **A.** Schematic showing the sequence and domain differences between the wild type FoxO1, the original reporter FT1, and the new FT2 reporter. Amino acid substitutions are listed below the protein map. Critical protein domains are highlighted: Forkhead DNA binding domain (FKH), nuclear localization sequence (NLS), nuclear exclusion sequence (NES), and the transactivation domain (TRANS). **B.** Schematic illustrating amino acids 311 to 330 of FoxO1 highlighting the phosphorylation sites (underlined) and the consensus phosphorylation sequence for the primary kinase that phosphorylates each site. Serine 326 and the reporter variants tested are highlighted in green. **C.** Time lapse images of HeLa cells stably expressing the respective reporters treated with IGF-I at 0 min and with PI-103 and LMB at 90 min. **D.** Average responses from a panel of HeLa cell lines stably expressing different FoxO1 reporter variants. Cell populations were treated with IGF-I at 0 min, and PI-103 and LMB at 90 min (*n* = 3 independent experiments per reporter, 100 cells per experiment). The sequence of amino acids of 325-327 of FoxO1 are listed to the right of the reporter template. The ‘3A’ refers to the reporter that has T24, S253, and S319 mutated to alanine. **E.** Rate of nuclear export following treatment with IGF-I for the different FT2 reporters graphed against the peptide charge. Black bar indicates the average response drawn from three independent experiments (shown as open circles). **F.** Measurement range (SFM nuclear localization to IGF-I nuclear localization) of individual HeLa cells stably expressing either the FT1 or the FT2-DDD constructs. Black bar indicates the population average of 500 cells (shown as open circles).

To construct a new reporter with increased specificity, we started from the template of the previously developed reporter, termed Fox Tracer 1 (FT1) and made alanine substitutions to 10 serine and threonine sites previously identified as phosphorylation sites of other kinases^23-27^ (Fig. 1A and Supplementary Fig. 1A, B). To reduce the protein size and eliminate other potential sources of non-specificity, we deleted a 234 amino acid segment from the C-terminus. Lastly, to increase the ease and efficiency of creating stable cell lines, we transferred the reporter from a lentiviral to a transposase-based genetic delivery system^28^ (Sup. Fig. 1C).

The first construct tested, FT2-LAP, had a significantly reduced variance and increased nuclear localization compared to FT1 under serum free conditions, which suggested an increase in specificity from the alanine mutations. However, this came at the cost of a reduced export rate following IGF-I treatment, which limits the ability to rapidly assess responses. We next tested reporter variants containing a series of mutations near S326, which is located in an AKT phosphorylation domain thought to regulate the rate of FoxO1 nuclear export by creation of a negatively charged region when phosphorylated^26, 29^ (Fig. 1B). The single mutation of alanine 326 back to serine (FT2-LSP) increased the rate of nuclear export, but also added back a DYRK1/Erk phosphorylation site^26, 29^(Fig. 1D, Sup. Fig. 1D). To eliminate this unwanted phosphorylation site, we mutated the peptide segment using the phosphomimetic amino acid aspartic acid. Furthermore, based on the concept that multiple negative charges better mimic the effect from phosphorylation^30, 31^, we tested four reporter variants that differed in the number of aspartic acid substitutions in this peptide segment. A comparison of these variants revealed that response rates increased with peptide charge, leading to a maximum rate of export for the FT2-DDD reporter (Fig. 1E, Movies 1-3).

As a control for the FT2-DDD reporter, we created a mutant with the three canonical AKT phosphorylation sites Thr24, Ser253 and Ser316 all mutated to alanine. As expected, this reporter (FT2-DDD 3A) does not translocate in response to IGF-I, which indicates that FT2-DDD responses to IGF-I were dependent on the canonical AKT phosphorylation sites (Fig. 1D). In summary, comparing the FT1 to the FT2-DDD reporter, we found that FT2-DDD is the optimal reporter for tracking the fidelity of IGF-I signaling responses based on its: increased specificity (Fig. 1A, B), reduced population variance in both serum free conditions and following IGF-I treatment (Sup. Fig. 1E), and increased measurement range (Fig. 1F).

### Responses to IGF-I stimulation are heterogeneous across the population

We next used the optimized FT2-DDD reporter to quantify responses of individual cells to IGF-I. As expected, we found a progressive increase in the average population response with increasing IGF-I dose, which was quantified as a change in the relative nuclear intensity of the fluorescent reporter in individual HeLa cells (Fig. 2A and B). Analysis of individual cells revealed heterogeneous responses to single doses of IGF-I and overlapping responses across multiple IGF-I doses, which has been interpreted as an inability of individual cells to discriminate different ligand doses^2^ (Fig. 2C). Importantly, this heterogeneity was not due to differences in the relative expression of the reporter between cells, nor differences in the basal level of activation in individual cells (Sup. Fig. 2). Thus, despite reducing possible effects of non-specificity in the readout for pathway activity, we find that responses to IGF-I remain heterogeneous across a population of individual cells.

**Fig. 2.**
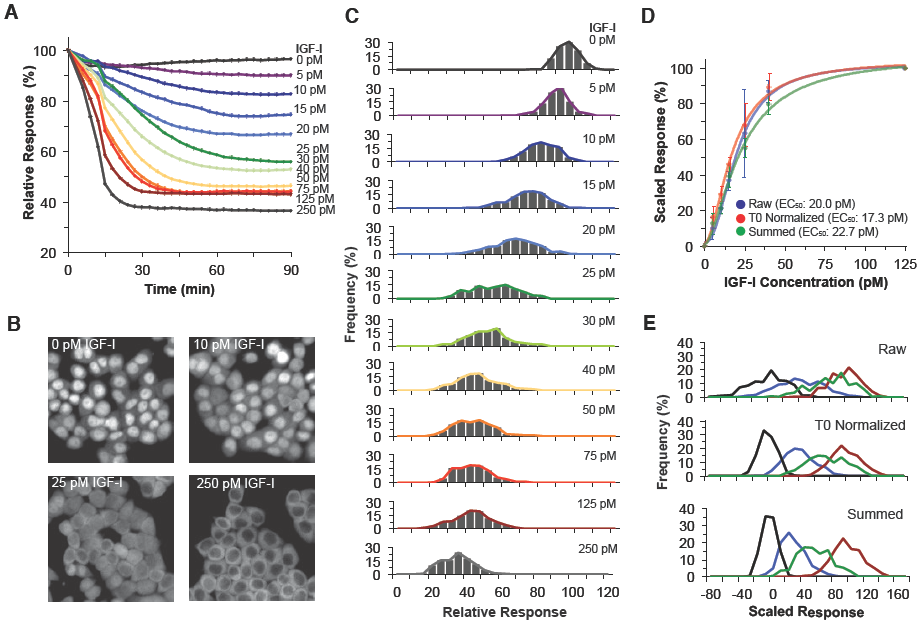
Reponses to IGF-I are heterogeneous across the population. **A.** Average responses of HeLa cells stably expressing the FT2-DDD reporter treated with different doses of IGF-I at 0 min. Doses are listed to the right in the respective order of the traces. (*n* = 200 cells per dose). Data have been normalized relative to the fluorescence intensity measured in SFM. **B.** Images of HeLa cells at 90 min following treatment with 0, 10, 25, or 250 pM IGF-I. Note the cell-cell variation in reporter localization after submaximal treatment with 10 or 25 pM. **C.** Frequency plots of individual cell responses to IGF-I at 90 minutes, color coded based on the IGF-I dose as in panel A (*n* = 200 cells per dose). **D.** IGF-I dose response curves calculated using either raw (blue), T0 normalized (red), or summed (green) measures. Error bars (standard deviation) and the average EC_50_ value are shown for each quantification method (*n* = 3 independent experiments). **E.** Comparison of individual cell responses to 0, 15, 25 or 125 pM IGF-I (color coded as in panels A and C) calculated using either raw, T0 normalized, or summed values (*n* = 200 cells per dose).

Integration of multiple time points to quantify a cell’s response has been shown to reduce the level of heterogeneity by reducing measurement noise^3^. To measure the effect of varying the number of measurements for each response, we compared dose-response relationships using either one image, which corresponds to the fluorescence at 90 min (Raw), two images, which allows cell responses at 90 min to be normalized based on their starting value (T0 normalized), or 15 images, which incorporates the signal change over a longer time window (Summed). We found that the EC_50_ values remained similar between measures (Fig. 2D), which indicates that the quantification method does not distort the relationship between doses. Yet, a comparison of the cell responses to 0, 15, 25, and 125 pM showed that overlap between the populations was reduced‐‐but not eliminated‐‐as the measurement window increased (Fig. 2D). These observations indicate that single cell responses to IGF-I are heterogeneous, even with use of analytical approaches that reduce variance.

### Responses of individual cells are stable across time

The Shannon-Hartley formula has been extensively used to quantify the signaling accuracy of biological systems^2-4^. A central assumption of the Shannon-Hartley channel capacity measurement, when applied to a population, is that the input-output relationship is identical for all cells in the population and observed variation is therefore attributed to noise in signal interpretation, transmission, and encoding. Based on this assumption, a noisy pathway would lead to varying responses in single cells over time (Fig. 3A, upper). Alternatively, there could be variation across the population in the input-output relationships of individual cells. In this case, we would expect to observe individual cells with minimal variation in signal transmission over time as well as heterogeneous responses across the population (Fig. 3A, lower). To separate these two scenarios, we quantified variation using the standard deviation of responses from each dose of IGF-I in the first 90-minute response window compared to the second 90-minute window when responses had reached a steady state (Fig. 3B). We found that the standard deviation during the first 90-minute response window increased as the dose of IGF-I increased (Fig. 3C, red line). This suggests different input-output relations across the cell population as seen by an increase in the number of responsive cells as the dose of IGF-I was increased. In contrast, the standard deviation for the second 90-minute window remained constant across doses and was similar to the 0 pM dose for the first 90-minute window (Fig. 3C, blue line). Assessment of the standard deviation calculated from the raw fluorescence measured in the first 90-minute window was similar across doses, which demonstrates that the increased measurement noise from unnormalized responses obscures the true population variation. (Sup. Fig. 3A). Together these observations indicate that individual cell responses are stable across time and that heterogeneity in the early response likely results from cells decoding the same IGF-I input signal into different FoxO1 signaling outputs.

**Fig. 3.**
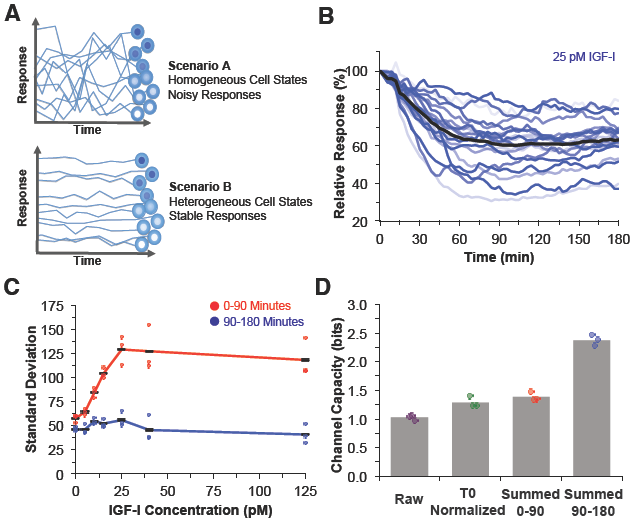
Single cell responses are heterogeneous across the population, but stable across time. **A.** Schematic of two scenarios where individual cell responses are heterogeneous because responses are noisy across time, or where cell responses are heterogeneous because cells exist in different states. **B.** Graph of single cell responses in different shades of blue following treatment with 25 pM IGF-I. The average response of the population is shown in black (*n* = 20 cells). **C.** Graph of the standard deviation from the dynamic response of cells treated with different IGF-I doses. The red trace with black bars shows the average standard deviation across 0-90 min and the blue trace shows the average standard deviation of the same cells from 90-180 min. Colored circles show the values from three independent experiments. **D.** Bar plot of the average channel capacity in bits using either raw, T0 normalized, or summed cell responses (bars) with the values from three independent experiments (colored circles).

Based on this interpretation, we hypothesized that variance quantified from stabilized responses provides a more robust calculation of noise for channel capacity measurements. To test this, we quantified channel capacity in various ways. Channel capacity calculated from a single time point yielded 1 bit (Fig. 3D), which is similar to prior measures of other signaling pathways^2^. This measure was increased to 1.2 bits when we computed channel capacity from the T0 normalized responses (Fig. 3D), and was further increased to 1.4 bits by using the summed responses drawn from 15 images. Finally, we quantified noise using the stable response window and found a channel capacity of 2.38 bits, which indicates that cells can discriminate at least 5 distinct IGF-I doses. While higher than previously reported, this is likely still an underestimate due to limitations in measuring individual cell responses. To test this idea, we treated cells with the nuclear export inhibitor Leptomycin B (LMB), which results in complete and sustained nuclear localization of the reporter. As a consequence, any variation in nuclear intensity after reaching steady state can be attributed to measurement noise rather than signaling noise. Comparison of how the signal varied across time for cells treated with LMB compared to cells treated with a submaximal dose of 17.5 pM IGF-I revealed nearly equivalent variances, which indicates that the true channel capacity may actually be much higher (Sup. Fig. 3B, C).

### Individual cells can recapitulate population-derived dose response curves to reveal high signaling fidelity

To test the capacity of a single cell to discriminate different signal intensities, we treated cells with multiple doses of IGF-I spread across the dose response range. Treatment with single doses of IGF-I served as a reference control (Fig. 4A). We found that the average population response to a single dose matched the response from cells treated with the same dose as part of a multiple-dose treatment (Fig. 4A), with some decrement in the signal when cells were maintained in IGF-I (e.g., see late phase of 125 pM response). Likewise, population frequencies for multiple-dose treated cells were nearly equivalent to time‐ and dose-matched controls maintained with a single dose (Fig. 4B). Further analysis of these multiple-dose data reveal that it is possible to recapitulate a dose response curve from individual cells, and moreover that this dose response is similar to a traditional population-level dose response curve in which different cell populations were treated with escalating doses of IGF-I (Fig. 4C).

**Fig. 4.**
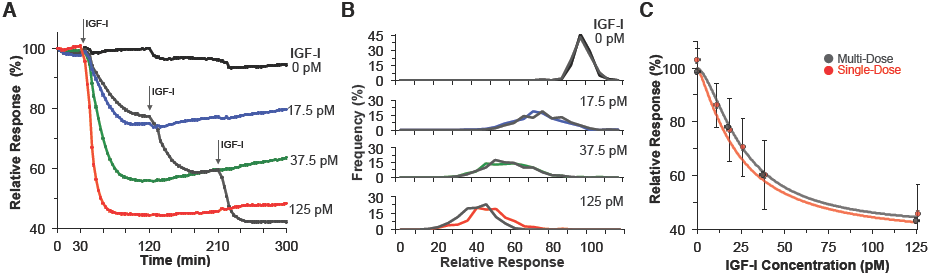
Population dose responses can be recapitulated in individual cells. **A.** Average responses of HeLa cells treated with 0 pM IGF-I 0-30min and then with either 17.5 pM (blue trace), 37.5 pM (green trace), or 125 pM (red trace) from 30-300 min. The gray trace shows cells treated with 0 pM IGF-I 0-30min, 17.5 pM 30-120 min, 37.5 pM 120-210 min and with 125 pM 210-300 min. The black trace shows the average response to cells maintained in 0 pM IGF-I. (*n* = 250 cells per condition) **B.** Frequency plots showing time and dose matched responses of cells exposed to a sustained treatment (colored lines as in panel A), or from multi-treated cells (gray lines). (*n* = 250 cells per condition) **C.** Graph of the average relative response against IGF-I dose. Red circles show average responses from different cell populations fit using logistic regression (red line) (*n* = 250 cells per dose). Error bars show the standard deviation from different populations treated with a single IGF-I dose. Gray circles show the average responses from the same cells treated with multiple IGF-I concentrations and fit using logistic regression (gray line).

We next sought to assess whether individual cells could discriminate multiple IGF-I doses and whether all cells have equivalent input-output relationships. We tracked 400 single cell responses after multi-dose treatment (0, 17.5, 37.5, and 125 pM IGF-I) and found that individual cells show distinct responses to each IGF-I dose (Fig. 5A, Movies 4-5). Moreover, we observed substantial variation across the population: some cells showed modest responses while others showed measurable responses to even the lowest dose tested. Comparison of traces from cells centered at the 25^th^ and 75^th^ percentiles revealed similar responses within the groupings but distinct differences between the groupings (Fig. 5B). Likewise, the average dose response relationships from these cell subpopulations and a subpopulation of cells centered at the 50^th^ percentile revealed consistent responses within the groupings (Fig. 5C). Moreover, a cell’s response to the first dose can be used to predict its response to the second dose (Sup. Fig. 4A). Finally, we find that responses of multi-dose treated cells are significantly different from time- and dose-matched cells maintained in the same dose (Sup. Fig. 4B). These studies indicate that individual cells can discriminate a minimum of four distinct IGF-I concentrations, and the input-output relationship is consistent in individual cells but varies across the population.

**Fig. 5.**
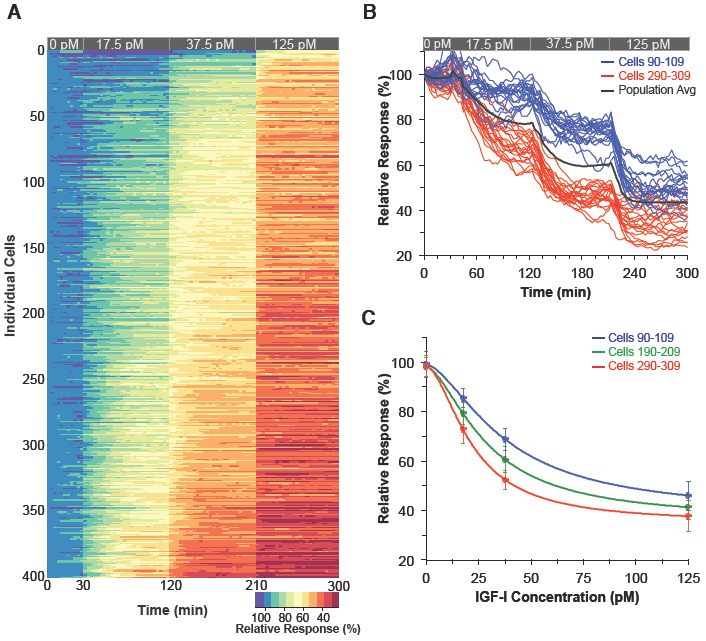
Accurate discrimination of multiple IGF-I doses in individual cells. **A.** Heat map showing individual HeLa cell responses across time to 0, 17.5, 37.5, and 125 pM IGF-I. Cells were sorted based on their summed response over 300 minutes. Graphed across time 0-30 min, 45-120 min, 135-210 min, and 228-300 min. (*n* = 400 cells). **B.** Single cell traces of 20 cells centered around the 75th (cells 90-109 blue traces) and 20 cells centered around the 25^th^ percentile (cells 290-309 red traces). The black trace shows the average response from the total cell population. **C.** Graph of the average relative response against IGF-I dose for cells 90-109 (blue), 190-209 (green), and 290-309 (red) fit using logistic regression (colored lines). Error bars show the standard deviation of each selected population.

### Single cell responses to PI3Kα inhibition are influenced by IGF-I activation level

Our multi-dose single cell data indicate that cells can discriminate multiple concentrations of IGF-I, and that a continuum of responses exists across the population due to different input-output relations. This raised the question of whether responses to pathway inhibition similarly vary across a population of cells. To test this, we treated cells with 25 pM IGF-I, a submaximal dose that leads to a wide variation in single cell responses (see Fig. 2B), followed by dose escalation of the PI3Kα inhibitor alpelisib (0-500 nM). At the population level, we observed the expected dose-response relationship to alpelisib treatment (Fig. 6A). To understand single cell responses to alpelisib, we compared cells from the upper (blue traces) and lower (red traces) quantiles after treatment with 25 pM IGF-I. We found that these two populations remain separable after inhibitor treatment (Fig. 6B, Movie 6), and that responses to PI3Kα inhibition were correlated with the level of activation from IGF-I (Fig. 6C). Comparison of all drug doses revealed similar separations between quantiles, up to 500 nM alpelisib, the dose at which signaling from IGF-I is nearly completely blocked (Fig. 6D). Together, these observations indicate that a cell’s response to inhibition is determined by its response to IGF-I, and as a consequence, responses to pathway inhibition are in line with a model in which there is variation in how cells encode a signaling input.

**Fig. 6.**
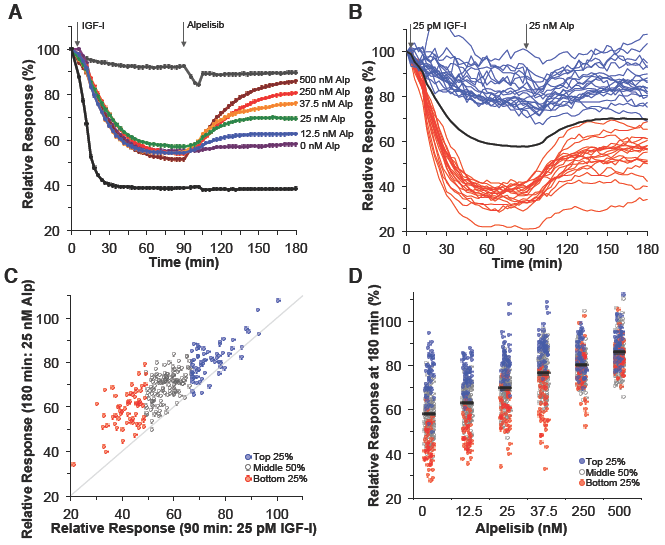
Single cell responses to PI3K inhibition are biased by the level of IGF-I activation. **A.** Time course of the average nuclear fluorescence intensities from HeLa cells maintained in SFM (light gray trace), treated with 125 pM IGF-I (dark gray), or treated with 25 pM IGF-I at time 0 min, and then at 90 min with 0 pM (purple), 12.5 nM (blue), 25 nM (green), 37.5 nM (orange), 250 nM (red), and 500 nM alpelisib (maroon). Average values are drawn from 250 cells. The first gray arrow indicates the time of IGF-I addition, and the second gray arrow indicates the time of alpelisib addition. **B.** Individual cell traces from 20 cells drawn from the top 25% (blue traces), and the bottom 25% (red traces) of cells treated at 0 min with 25 pM IGF-I and at 90 min with 25 nM alpelisib. The black trace shows the average response of the total population across time. **C.** Plot of individual cell responses at 90 min after IGF-I treatment against responses after 25 pM alpelisib (180 min). Cells are color coded based on their response at 90 min. Dotted gray line shows a 1:1 relationship between the 90 and 180 min responses. (*n* = 250 cells) **D.** Plot of the relative responses of individual cells at 180 min drawn from panel A. Individual cell responses are color coded such that blue circles show the top 25% of cells at 90 min, gray open circles show the middle 50% of cells at 90 min, and the red circles show the bottom 25% of cells at 90 min. The average population response at 180 min is shown with the black bar. Responses from 250 cells are plotted per treatment.

## Discussion

Understanding how individual cells respond to diverse stimuli is a fundamental issue in cell biology^32-34^. To that end, we describe here the development of a FoxO1 reporter with a higher specificity for IGF-I signaling compared to our previous FoxO1 reporter^10^. With this improved reporter, we observed that individual cell responses to IGF-I still overlapped between doses, suggesting low signaling fidelity. Further analysis showed that this overlap was from differences in how cells encode the signal, rather than from noise in the signaling pathway. Multi-dose treatment protocols validated these findings and showed that cells could accurately discriminate at least four different IGF-I doses. These studies demonstrate that individual cells can accurately process extracellular growth factors into downstream signaling responses despite the observation that population responses overlap between doses.

### Reporter optimization for live cell imaging studies

Genetically encoded fluorescent reporters provide a powerful tool for understanding and quantifying dynamic signaling responses^35^. Various types of signaling reporters have been developed, including FRET sensors^36, 37^, synthetic translocation reporters^38^, and native proteins that translocate between the nucleus and cytoplasm (FoxO1^10, 39^, p53^40^, SMAD2^41^, NFkB^42^, Erk^43^). Native proteins are advantageous in that they are biologically tuned to respond under physiological conditions and the function of critical protein regions are typically well-described. Here we used knowledge of the FoxO1 sequence to optimize an existing FoxO1 reporter. Specifically, we increased specificity by both targeting a large peptide region for deletion that is not involved in nuclear translocation and by mutating multiple phosphorylation sites that are not phosphorylated by AKT. Additionally, we optimized nuclear export through the modification of a short peptide sequence. These modifications would have been difficult to predict in the absence of previous studies. For example, nuclear export was improved by substitution of a phosphomimetic amino acid rather than an alanine at serine 326, and export was further aided by mutating the surrounding amino acids to better match the charge from phosphorylation^30, 31^. All together these modifications significantly increased the specificity of the reporter for IGF-I signaling in HeLa cells, and allowed for increased accuracy in tracking individual cell responses. Our approach to optimize the FoxO1 reporter, which considers the molecular structure and function of FoxO1, could be used to efficiently develop reporters of other signaling pathways.

### Channel capacity quantification

Previous studies of the Erk, NFkB, and Ca^2+^ signaling pathways have found channel capacities from 1-1.7 bits^2, 3, 5, 44^. Here, we assessed the channel capacity of the PI3K-AKT pathway using IGF-I. Importantly, this growth factor produces sustained Akt signaling responses^45^, making it simpler to quantify and identify signaling patterns across time as compared to transient responses. We found that using a single time point to quantify channel capacity yielded a value of 1 bit, which was increased to 1.4 bits by normalization and incorporation of measures from multiple time points. These values are in line with previous measures on other signaling pathways^2, 3, 5, 44^. We further refined our calculations by considering that single cell responses are sustained over time and do not show large fluctuations. Thus, rather than using the heterogeneity in the extent of responses as has been previously used in prior applications of the Shannon-Hartley formula to quantify signaling responses, we used the variance across time once cells had reached a steady state in their response. This new approach that posits that each cell has its own reliable communication channel, rather than the population having a single noisy communication channel, increased the calculated channel capacity to 2.38 bits (∼5 doses), nearly double previous calculations for other pathways. Based on these results, we hypothesize that other signaling pathways may also have a much higher signaling fidelity then previously reported.

### Multi-treatment experiments reveal variation in single cell responses

The ability to quantify responses to multiple treatments in individual cells provides a powerful technique for understanding mechanisms of population variation^46, 47^. This is in stark contrast to standard experiments in which single measures on individual cells are drawn from different populations, or the response to only a single stimulus is tracked across time. We used this multi-treatment approach to achieve several means: first we used LMB normalization to quantify the measurement range of the reporter and to show that individual cell responses are not biased by the level of reporter expression, second we used multi-dose IGF-I treatments to validate that cells can accurately discriminate multiple doses of IGF-I and to identify variability in IGF-I encoding across the population, and third we used activation-inhibition experiments to determine that the level of signaling activity in an individual cell biases the extent of inhibition. Without multiple measurements on the same cells, it would not be possible to identify these patterns nor accurately quantify potential sources of biological variation.

### Cells exist in different states with different input-output relationships

Our data indicate that cells can accurately interpret multiple growth factor signals in their extracellular environment and that across the population there is variation in how cells encode this signal. As a consequence, it is not possible to accurately decode the input based on a single response. It is only through knowledge about how a cell responds to a second perturbation—either activation or inhibition—that the input signal could be decoded. This raises the question of what cellular parameter varies to create different input-output relationships across the population. It is reasonable to assume that variation in the input-output relationships between cells occurs because they differ in the content of the IGF-I receptor or another pathway component^48^, but it could also be from differences in basal signaling^49^, cell shape^50^, nuclear import/export machinery^38^, or phosphatase activity. Assessing each of these mechanisms independently is challenging, but our work suggests that it may be possible to tease apart these different mechanisms using a single-cell multi-treatment approach.

Our studies indicate that signaling responses to molecules that activate or inhibit the IGF-I-AKT signaling pathway are not necessarily noisy. This is important as it rules out one mechanism for drug resistance in cancer: that cells have large and variable signaling fluctuations that allow them to overcome inhibition. This conclusion also suggests a role for microenvironmental signals to modulate therapeutic response. For instance, subpopulations of cells may be subject to signals that lead to pathway activation, whereby a larger drug dose may be required to fully inhibit these cells^19, 51^. These experiments open the door to more in depth studies on how various pathway inhibitors alter signaling activity and how treatments regimens could be optimized for use in the clinic, particularly in the metastatic setting where cancer cells have disseminated to distinct tissue sites enriched with unique complements of signaling molecules.

## Summary

A significant question in cell biology is how much noise is present in a signaling pathway, and as a consequence how accurately a signal can be interpreted and processed. Along with the development of an improved reporter, we found that individual cells can faithfully interpret multiple IGF-I concentrations, although each cell encodes the signal differently. This conclusion has significant impacts on how we think about the regulation of biochemical pathways and cell-cell heterogeneity.

## METHODS

### Cloning

The new FoxO1-Clover reporter was constructed from a modified pSBbi-RP plasmid^28^ (plasmid number: 60513 Addgene, Cambridge, MA) that had dTomato replaced with mCherry appended with N‐ and C‐ terminal nuclear localization signal (NLS) tags. To recreate the FoxO1-Clover reporter using the new transposase vector, pLenti-FoxO1-Clover^10^ was digested using NaeI and XbaI restriction sites and ligated into the modified pSBbi plasmid. To create the FT2-LAP reporter, DNA for FT2-LAP was synthesized (IDTDNA) with the following amino acid substitutions: S209A, H212R, S215A, S246A, S284A, S295A, S298A, S300A, S326A, S380A, S391A, T399A. The synthesized DNA contained 5’ BglII and 3’ Hind3 restriction sites that enabled ligation into the pSBbi-FT1 plasmid. Mutations to FoxO1 amino acids 325-327 (LSP peptide) were introduced via PCR using primers containing the desired mutation/s and ligated in frame using ScaI and Hind3 restriction enzyme sites. DNA for the FT2-DDD-3A construct was synthesized (IDTDNA) and contained six additional substitutions compared to FT2-LAP: T24A, S253A, S316A, L325D, S325D, and P327D. The FT2-DDD (plasmid number: 106278) and the FT2-DDD-3A (plasmid number: 106279) are available from Addgene.

### Transfection and selection of stable cell lines

HeLa cells, validated by STR profiling, were grown in DMEM supplemented with 10% FBS. To create stable lines expressing each reporter variant, cells were transfected using Lipofectamine 3000 (ThermoFisher) with the respective FoxO1-clover expression plasmid and the transposase expression plasmid pSB100X (plasmid number: 34879, Addgene, Cambridge, MA) at a ratio of 4:1. Two days following transfection cells were selected by incubation with puromycin (2 μg/ml) for 7-days. Single cell clones expressing FT2-DDD were created through limiting dilution in 96-well plates.

### Microscopy

Live cell imaging was performed using an EVOS FL Auto microscope (ThermoFisher) with a stage top incubator maintained at 37°C, and in 95% air, 5% CO2. Cells were imaged every 3 minutes using a 10X fluorite objective (N.A. 0.3), and a GFP LED light cube (excitation peak, 472/22 nm; emission peak, 510/42 nm). Images were pre-processed using the ImageJ plug-ins (NIH, Bethesda, MD) StackReg (image translation) and Gaussian Blur (2 pixels). Background fluorescence was calculated per plate for each time point by taking the average fluorescence of three regions of interest that did not contain cells. To quantify signaling responses, individual cells were manually tracked by selecting a location in the center of each cell nuclei using the ROI plug-in. The fluorescence intensity of that location was then assessed across time. Cells that migrated over the course of the experiment such that the T0 location no longer aligned with later time points were excluded from analysis. Additionally, cells that divided, died, were multi-nucleated, or overlapped with another cell were excluded from analysis.

Three measures were used to quantify individual cell responses: the ‘nuclear localization’, the ‘relative nuclear intensity’, and the ‘scaled relative nuclear intensity’. The ‘nuclear localization’ for each cell was calculated by normalizing fluorescence intensities to the nuclear fluorescence intensity 90 min after leptomycin (LMB) treatment. The ‘relative nuclear intensity’ for each cell was calculated by normalizing fluorescence intensities to the nuclear fluorescence intensity at T0 following 90 min in serum free media. The ‘scaled relative nuclear intensity’ was calculated by scaling each cell’s response to the population average, where 0 pM was set to 0 and 125 pM IGF-I was set to 100.

### Reporter Characterization

To characterize each HeLa reporter cell line, cells were serum starved for 90 minutes in Fluorobrite media (ThermoFisher) supplemented with 2 mM L-glutamine, 0.1% bovine serum albumin, penicillin and streptomycin. Cells were then treated with 250 pM R3-IGF-I for 90 minutes followed by the combined addition of the dual PI3K and mTOR inhibitor PI-103 (1 μM) and LMB (100 nM) for 90 minutes. From these experiments, we quantified: the average rate of reporter nuclear exclusion following IGF-I treatment, the average rate of nuclear localization following PI-103 and LMB, and the average measurement range, which was defined as the difference between the percent nuclear localization after serum starvation and the percent nuclear localization 90 min after IGF-I treatment. Export and import rates were calculated using the population average response and fit via non-linear regression (Graph Pad Prism 7). All cell lines were imaged using the same 75 millisecond imaging duration.

### IGF-I single and multi-dose experiments

For IGF-I dose response experiments, HeLa cells were plated in 12-well plates (Falcon) allowed to grow for 48 h and then serum starved for 90 min in imaging media. Cells were then treated with: 0, 5, 10, 15, 20, 25, 30, 40 50, 75, 125, or 250 pM R3-IGF-I. For multi-dose experiments, cells were imaged 30 minutes prior to IGF-I addition and then treated with 0, 17.5, 37.5, or 125 pM R3-IGF-I for 270 min. In separate wells, cells were imaged in SFM for 30 minutes and then treated with three progressively higher doses of IGF-I for 90 min each (17.5, 37.5, or 125 pM IGF-I). Dose response curves were fit via non-linear regression using the Hill equation (Graph Pad Prism 7). ‘Raw’ responses were quantified using the nuclear intensity 90 min after IGF-I treatment. ‘T0 normalized responses’ were calculated by taking the 90 min value relative to the 0 min value. ‘Summed responses’ were calculating by summing the relative change of the T0 normalized data using 15 equally spaced images (every 6 min). Individual cell responses to multiple doses of IGF-I were sorted using the summed responses across the 300-minute experiment. Heat maps were created in R (https://www.R-project.org) using the gplots heatmap package version 3.0.1.

### Leptomycin B experiments

To quantify the relationship between reporter expression and sub-maximal IGF-I responses, cells were serum starved for 90 minutes and then treated with 17.5 pM R3-IGF-I. After 90 minutes cells were treated with 1 μM PI-103 and 100 nM LMB for 90 minutes. To track variation in individual responses across time, cells were treated with either 100 nM LMB, or 17.5 pM R3-IGF-I for 180 minutes.

### Alpelisib experiments.

For tracking cell responses to PI3Kα inhibition, cells were serum starved for 90 min and then treated with either 0, 25, or 125 pM R3-IGF-I. After treatment for 90 min cells were exposed to 0, 12.5, 25, 37.5, 250, or 500 nM alpelisib for 90 min.

### Quantification of channel capacity

Channel capacity was calculated using the Shannon-Hartley formula: CC = ½ log_2_(S^2^/N^2^), where CC is the channel capacity in bits, S is the signal magnitude quantified by taking the variance of the average response across six relatively evenly spaced IGF-I doses representing the full dose range (0, 10, 15, 25, 40, 125 pM R3-IGF-I), and N is the noise magnitude quantified by taking the average variance within a dose and averaging across all doses^3^.

### Data Availability

Source data are available from the corresponding author upon reasonable request.

## Acknowledgements

These studies were supported by NIH research grants to 1U54CA209988 (L.M.H.), U54-HG008100 (L.M.H.) and the Prospect Creek Foundation (L.M.H.). L.M.H. was also supported by the Jayne Koskinas Ted Giovanis Foundation for Health and Policy and the Breast Cancer Research Foundation, private foundations committed to critical funding of cancer research. The opinions, findings, conclusions or recommendations expressed in this material are those of the authors and not necessarily those of the Jayne Koskinas Ted Giovanis Foundation for Health and Policy or the Breast Cancer Research Foundation or their respective directors, officers, or staffs.

## Authors’ contributions

S.M.G. designed the study, performed experiments, and analyzed data; M.A.D and E.B. analyzed data. L.M.H. supervised the study. S.M.G. and L.M.H. wrote the manuscript. All authors participated in editing the manuscript.

---

**Movie 1 (connects to Figure 1).** Tracking of FT1-clover in HeLa cells during exposure to 250 pM IGF-I at 0 min followed by exposure to 1μM PI-103 and 100 nM LMB at 90 min. Images were collected every 3 min for 180 min by time-lapse epi-fluorescence microscopy (Evos FL Auto microscope). The video playback rate is 5 frames per second.

**Movie 2 (connects to Figure 1).** Tracking of FT2-LAP-clover in HeLa cells during exposure to 250 pM IGF-I at 0 min followed by exposure to 1μM PI-103 and 100 nM LMB at 90 min. Images were collected every 3 min for 180 min by time-lapse epi-fluorescence microscopy (Evos FL Auto microscope). The video playback rate is 5 frames per second.

**Movie 3 (connects to Figure 1).** Tracking of FT2-DDD-clover in HeLa cells during exposure to 250 pM IGF-I at 0 min followed by exposure to 1μM PI-103 and 100 nM LMB at 90 min. Images were collected every 3 min for 180 min by time-lapse epi-fluorescence microscopy (Evos FL Auto microscope). The video playback rate is 5 frames per second.

**Movie 4 (connects to Figure 5).** Tracking of FT2-DDD-clover in HeLa cells during exposure to 0 pM IGF-I at 0 min followed by exposure to 37.5 pM IGF-I at 30 min. Images were collected every 3 min for 300 min by time-lapse epi-fluorescence microscopy (Evos FL Auto microscope). The video playback rate is 5 frames per second.

**Movie 5 (connects to Figure 5).** Tracking of FT2-DDD-clover in HeLa cells during exposure to 0 pM IGF-I at 0 min followed by exposure to 17.5 pM IGF-I at 30 min, 37.5 pM IGF-I at 120 min and 125 pM IGF-I at 210 min. Images were collected every 3 min for 300 min by time-lapse epifluorescence microscopy (Evos FL Auto microscope). The video playback rate is 5 frames per second.

**Movie 6 (connects to Figure 6).** Tracking of FT2-DDD-clover in HeLa cells during exposure to 25 pM IGF-I at 0 min followed by exposure to 25nM Alpelisib at 90 min. Images were collected every 3 min for 180 min by time-lapse epi-fluorescence microscopy (Evos FL Auto microscope). The video playback rate is 5 frames per second.

**Supplementary Figure 1. FT2 Reporter creation and characterization. A.** Table of the different reporters tested, the length of FoxO1 in amino acids (aa) for each reporter, the number of mutations, and a list of the specific mutations. **B.** List of phosphorylation sites in FoxO1 that were mutated in FT2, and the surrounding 15 amino acids (phosphorylation site highlighted in red). **C.** Schematic of the genetic segment inserted into the genome via the transposase. The bi-directional blue arrow shows the RPBSA promoter (light blue) that drives expression of mCherry appended with nuclear localization sequences (NLS) on the N‐ and C-termini, followed by the T2A cleavage sequence, and puromycin resistance (Puro). The other promoter is EF1a which drives expression of the FT2-clover protein. **D.** Table showing a list of the reporters tested in HeLa cells and the average IGF-I nuclear export rate, PI-103/LMB nuclear import rate, percent localization of the reporter in serum free media (SFM) relative to the value from PI-103/LMB, and the percent localization of the reporter after treatment with 250 nM IGF-I relative to the value from PI-103/LMB. Averages are from the mean value of three independent experiments +/− the standard deviation. **E.** Frequency plots of FT1 HeLa cells in SFM relative to LMB (solid black line) and after treatment with 250 nM IGF-I (dotted black line) plotted with FT2-DDD HeLa cells in SFM relative to LMB (solid blue line) and after treatment with 250 nM IGF-I (dotted blue line). (*n* = 500 cells per condition).

**Supplementary Figure 2. Single cell responses to IGF-I are heterogeneous across the population. A.** Population average (*n* = 250 cells) from a monoclonal population HeLa cells treated at time 0 with 17.5 pM IGF-I and then at 90 min with 1μM PI-103 and 100 nM LMB. The standard deviation is shown in blue. **B.** Dot plot showing the raw fluorescence of individual cells after LMB/PI-103 treatment against the fluorescence in SFM drawn from the mean plotted in panel (*n* = 500 cells). **C.** Dot plot of a multiclonal HeLa line (red dots) graphed with a monoclonal population of HeLa cells (blue dots) each expressing the FT2-DDD reporter. (*n* = 500 cells per reporter). **D, E, F.** Dot plots of the same cells shown in A and B with LMB fluorescence graphed against the relative response to 17.5 pM IGF-I as assessed using the SFM relative response (D), the LMB normalized response (E), or the summed response (F). (*n* = 500 cells).

**Supplementary Figure 3. Single cell responses to IGF-I are stable across time. A.** Graph of the average standard deviation (black bars) from the raw fluorescence at 90 min after treatment with different concentrations of IGF-I. The blue circles show the standard deviation from each of three independent experiments. **B.** Single cell traces of HeLa cells treated at time 0 with 100 nM LMB (green traces) or 17.5 pM IGF-I (blue traces). **C.** Frequency plot showing that mean signaling deviation from 90 to 180 min following treatment with either 17.5 pM IGF-I (blue) and 100 nM LMB (green). The standard deviation of the population response for each treatment is shown in the upper right corner. (*n* = 500 cells per condition).

**Supplementary Figure 4. Accurate discrimination of multiple IGF-I doses in individual cells. A.** Plot of single cell responses at 120 min compared to responses at 210 min from cells maintained in the same dose (sustained: black circles) or treated with 37.5 pM IGF-I (treated: red circles). Responses were linearly fit (dotted black line) and the coefficient of determination (R^2^) is shown for each population. **B**. Plot of the summed responses from 250 individual cells treated with sequential IGF-I treatments from 30-120 min, 120-210, and 210-300 min compared against control populations where the comparable IGF-I dose was maintained (*n* = 250 cells per condition). Black bars show the population average. Differences between sustained and treated responses for the three time periods were compared using unpaired t-tests and yielded P values that were all less than 0.0001.

